# Viral Strategies for In Vivo Optogenetic Stimulation of Purkinje Cell Terminals and Electrophysiological Characterization of Downstream Cerebellar Nuclear Activity in Mice

**DOI:** 10.64898/2026.09.27.754842

**Authors:** Esteban Merino, Se Jung Jung, Evangelos G. Antzoulatos, Diasynou Fioravante

## Abstract

The cerebellar cortex regulates motor and cognitive function through its sole output neurons, the Purkinje cells (PCs), which inhibit downstream targets, yet precise manipulation of PC activity in vivo remains technically challenging. Here, we describe two viral strategies with differing cell-type specificities for in vivo optogenetic control of PCs and functional assessment of neural activity in downstream target regions. Specifically, we detail strategies for expression of the fast channelrhodopsin ChETA in PCs that enable light-evoked activation of PC axonal terminals in medial deep cerebellar nuclei (mDCN), and demonstrate light-evoked modulation of mDCN neural activity using in vivo extracellular electrophysiology. This protocol provides a reproducible and adaptable framework for cell-type-specific manipulation and functional characterization of cerebellar circuits, with broad applicability to studies of cerebellar physiology, motor and nonmotor behavior, learning and disease.

**SUMMARY:** This protocol describes viral strategies for in vivo optogenetic tagging of cerebellar Purkinje cells and extracellular electrophysiological recording of opsin-evoked responses in the medial deep cerebellar nucleus, providing a reproducible approach to investigate cerebellar circuit connectivity and functional interactions with cell-type specificity.

## INTRODUCTION

The cerebellum (CB) regulates motor coordination and learning, as well as cognitive, affective, and social behaviors, through the integration of diverse incoming information and the generation and propagation of prediction and prediction-updating signals that influence neural computations in downstream brain regions^1–5^. This input-output transformation depends critically on Purkinje cells (PCs), the sole output neurons of the cerebellar cortex, which provide powerful inhibitory input principally to the deep cerebellar nuclei (DCN), as well as to brainstem structures including the vestibular and parabrachial nuclei^6–9^. DCN neurons integrate PC inhibition with excitatory and neuromodulatory inputs arising outside the CB to generate the principal excitatory CB output^10^. Thus, understanding how CB cortical activity shapes CB output requires determining how PC activity regulates DCN neuronal firing in vivo.

The anatomical organization and intrinsic and synaptic electrophysiological properties of PCs are well characterized^11–19^, providing a strong foundation for investigating how CB cortical activity shapes DCN output. However, PCs are not a homogeneous population: their molecular identity, intrinsic firing properties, and physiological responses vary across CB regions (e.g., vermis, flocculus)^9,20^. PC projections to the DCN are also topographically organized, with distinct PC populations innervating different nuclei (medial, interposed, and lateral DCN) and subnuclear territories^10^. Consequently, resolving the pathway- and territory-specific influence of PCs on CB output requires selective control of anatomically defined PC subsets combined with electrophysiological recordings of DCN activity in vivo.

In the CB, viral approaches offer experimental flexibility by enabling anatomically restricted opsin expression without requiring transgenic animals, with applicability across species^21–23^. By exploiting the unique position of PCs as the sole projection neurons of the CB cortex, localized viral delivery to specific CB cortical regions enables tracing and manipulation of spatially defined PC populations and their projections to the DCN. Alternatively, the topographic organization of PC projections to the DCN can be exploited through retrograde viral delivery to selectively target PC populations innervating defined nuclear territories. Together, these complementary strategies enable anatomically restricted manipulation of PC activity and investigation of its functional influence on CB output.

Successful optogenetic manipulation of the PC-DCN circuit requires consideration of the physiological properties of both presynaptic and postsynaptic neurons. PCs exhibit high spontaneous firing rates, which can transiently exceed 100 Hz in behaving animals^24–26^. Moreover, PCs provide tonic inhibition to DCN neurons, which are intrinsically active despite receiving converging inhibitory input from multiple PCs^27,28^. Optogenetic stimulation must therefore recruit sufficient PC inhibitory input to reliably suppress ongoing DCN firing while avoiding stimulation regimes that perturb PC axonal excitability or evoke post-inhibitory rebound activity in DCN^29–31^. Achieving this requires careful selection of opsins, robust expression and functional localization at PC axonal terminals, and optimization of optical stimulation parameters. For channelrhodopsin specifically, PC axons require longer light pulses than PC somata to fire reliably and are more susceptible to light-induced perturbation^31^, so parameters cannot be adopted directly from somatic stimulation regimes. Combining localized optogenetic stimulation of PC terminals with simultaneous in vivo electrophysiological recordings of DCN activity is a powerful approach for establishing stimulation parameters that produce reliable inhibition while minimizing unintended physiological responses. Previous studies have demonstrated that optogenetic stimulation of PC axonal terminals within the DCN can modulate DCN neuronal firing and influence CB-dependent behaviors^32–35^. These studies used genetic approaches that drive opsin expression broadly across PCs, providing extensive PC coverage well suited to questions about CB cortical output as a whole. Questions concerning specific PC populations instead require expression restricted to a defined cortical region or projection target, which viral delivery provides.

Here, we present two complementary adeno-associated virus (AAV)-based strategies for anatomically restricted optogenetic control of PC populations using the fast channelrhodopsin ChETA^36^, enabling selective activation of PC axonal terminals in mDCN. We describe the experimental workflow, from stereotaxic viral delivery and histological validation of opsin expression to localized optical stimulation, simultaneous in vivo extracellular recordings, and single-unit analysis. We identify stimulation parameters that produce robust inhibition of mDCN neuronal firing without significant post-inhibitory rebound firing loss of inhibitory efficacy suggestive of PC axonal depolarization block. This protocol provides an adaptable framework for investigating how anatomically defined PC populations regulate CB output across different CB regions and animal models.

## PROTOCOL

Adult (6-8 weeks old) C57BL/6J or Ai14 (B6.Cg-Gt(ROSA)26Sor^tm^^14(CAG-tdTomato)Hze^/J, The Jackson Laboratory #007914) mice were housed under controlled environmental conditions (22 ± 1 °C; 12-h light/dark cycle) with ad libitum access to food and water. All experimental procedures were approved by the Institutional Animal Care and Use Committee (IACUC) of the University of California, Davis, and were conducted in accordance with the National Institutes of Health (NIH) Guide for the Care and Use of Laboratory Animals.

### 1. Pre-surgical preparation and anesthesia

**1.1.** Sterilize all surgical instruments (**Fig. 1**) by autoclaving before surgery. Thoroughly disinfect the surgical area with 70% ethanol.

**Figure 1:**
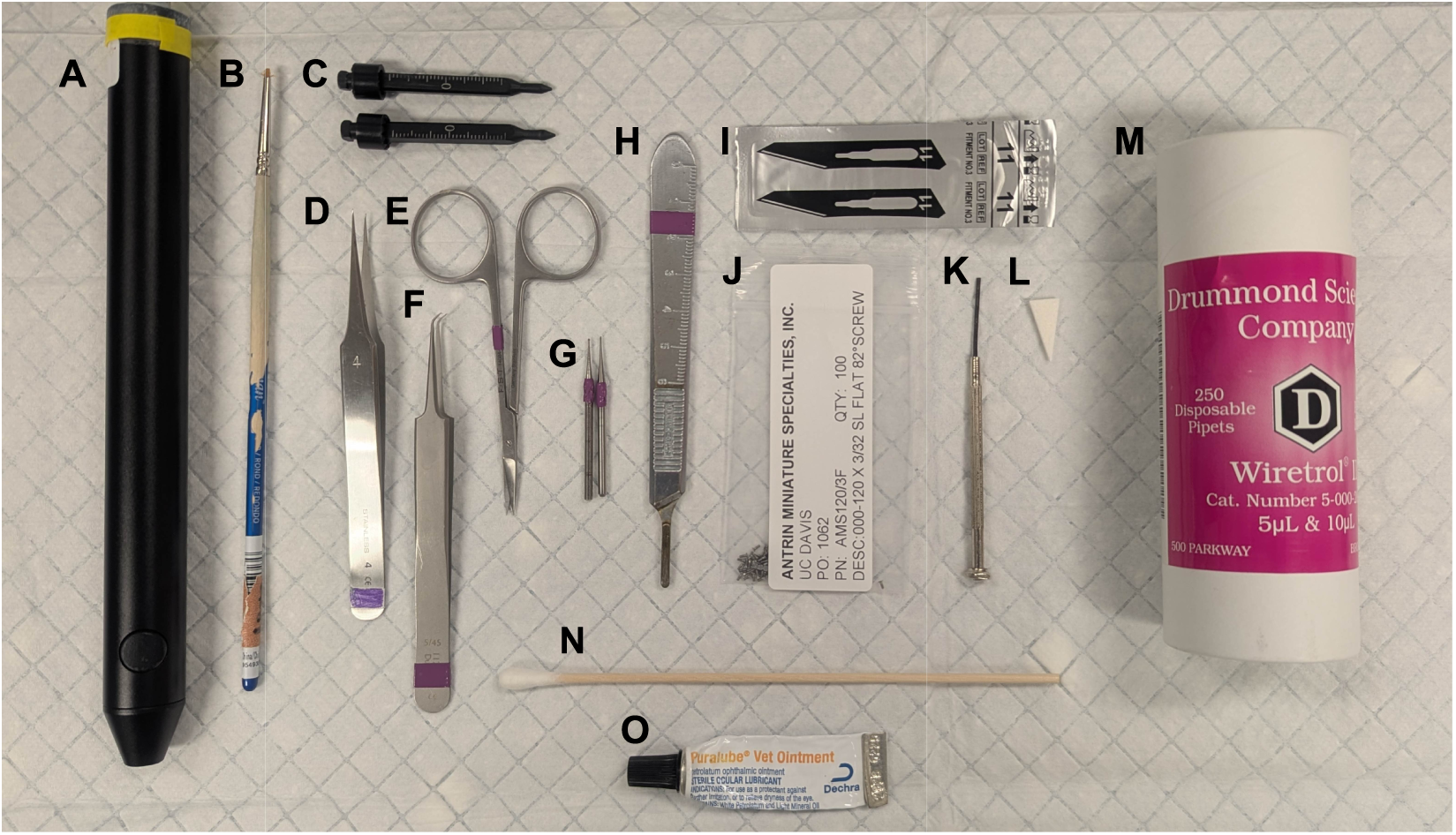
Surgical instruments and equipment used in experimental procedures. Cordless microdrill (**A**) and carbide burs (**G**) used for craniotomy and preparation of the skull for probe implantation; Cotman Watercolour Brush Series 111, No. 0, with shortened bristles (**B**), used to apply fluorescent dextran to the probe; Mouse non-rupture ear bars (**C**) used to secure and stabilize the animal during surgery; Dumont #4 forceps (**D**) and Bonn scissors (**E**) used for tissue manipulation and skin incision, respectively; Dumont #5/45 forceps (**F**) used for precise manipulation of tissue and surgical materials; Bard-Parker #3 stainless steel surgical blade handle (**H**) and sterile #11 surgical blades (**I**) used for skin incision; Miniature flat-head machine screws (**J**) used as the skull ground/reference electrode; a screw is implanted in contact with the brain surface and connected by a wire to the probe reference; Flat-head slotted screwdriver (**K**) used to place and secure the miniature skull screws; Sterile absorption triangles (**L**) used to absorb blood and excess fluids during surgery; Drummond Wiretrol II calibrated glass pipettes (**M**), pulled to a fine tip with a pipette puller to administer viral vectors and to mark trephination sites on the skull; Cotton-tipped dental applicators (**N**) used for cleaning, drying, and applying materials during surgery; Puralube ophthalmic ointment (**O**) applied to protect the eyes from drying during anesthesia. For detailed information on the materials and equipment used in the surgical procedures, please refer to the Materials List provided with this article.

**1.2.** Using a pipette puller, pull glass micropipettes to a fine tip (10-20 µm in diameter). Back-fill a pipette with 1% Fast Green solution and mount it onto a stereotaxic microinjection system. This pipette will be used to identify stereotaxic landmarks on the skull.

**1.3.** Transfer mice to the surgical room in covered home cages to minimize stress before anesthesia.

**1.4.** Weigh each mouse to calculate the required drug dosages.

**1.5.** Place the mouse in an induction chamber and anesthetize with isoflurane (5% for induction, 1–2% for maintenance; **CAUTION:** Isoflurane is a volatile anesthetic. Perform all procedures in a properly ventilated area or under a scavenging system to minimize occupational exposure) delivered in oxygen (0.8–1.0 L/min).

**1.6.** Once the mouse reaches surgical-plane anesthesia, defined as lack of paw withdrawal response to toe pinch, transfer to a stereotaxic frame positioned on a feedback-controlled heating pad to maintain body temperature throughout the procedure (Fig. 2A).

**Figure 2:**
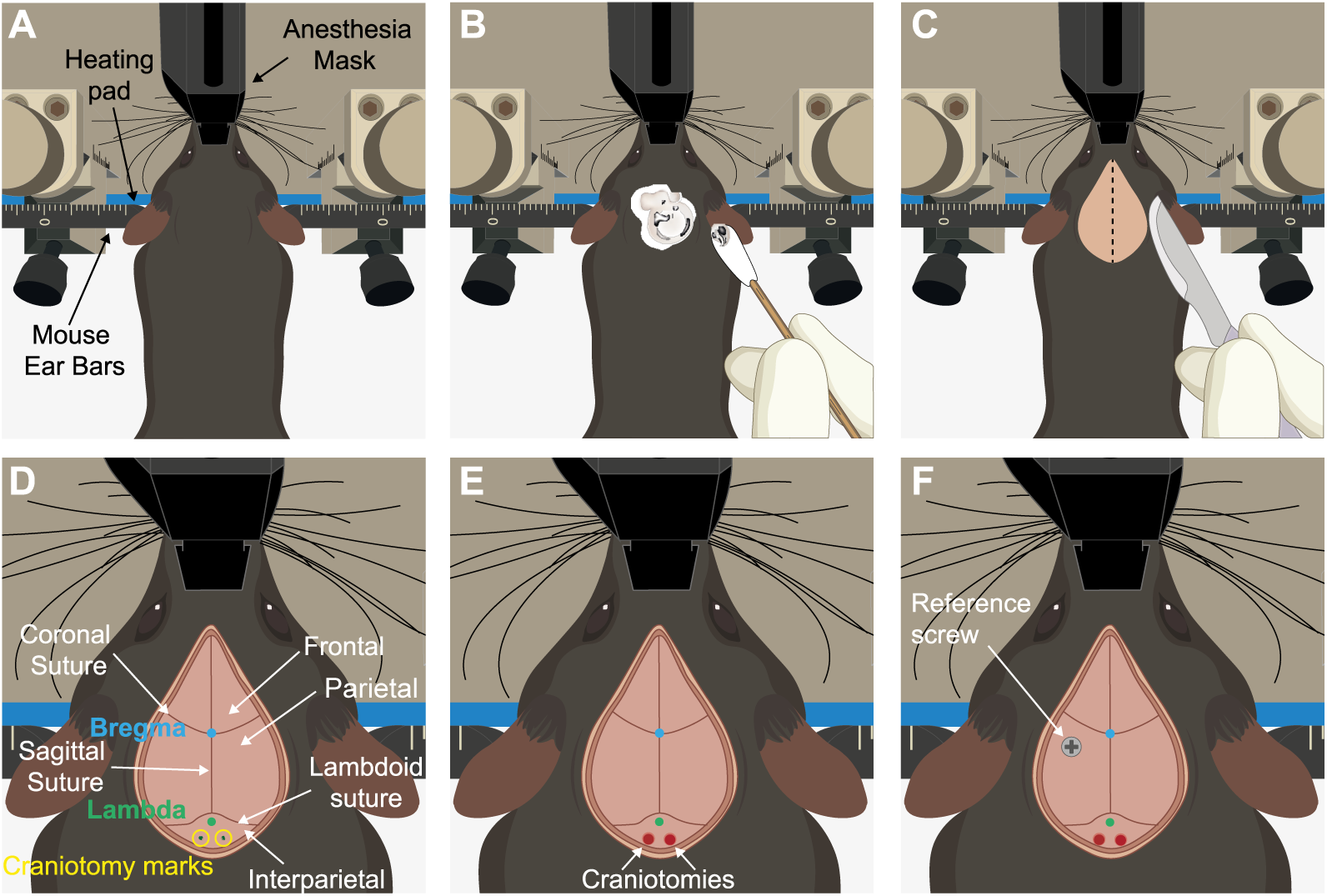
Surgical preparation and cranial access for viral injection and in vivo electrophysiological recordings. All images are shown in dorsal view. **A**) Anesthetized animal properly positioned in the stereotaxic frame on a feedback-controlled heating pad. **B**) Application of depilatory cream to remove fur from the head region. **C**) Surgical field after fur removal, with a dashed line indicating the location of the midline incision used to expose the skull. **D**) Exposed skull following skin incision, showing the main anatomical landmarks used for stereotaxic positioning, including the coronal, sagittal, and lambdoid sutures, as well as bregma and lambda. The frontal, parietal, and interparietal bones and the approximate locations of the craniotomies are also indicated. **E**) View of the skull after craniotomies for viral injection and/or electrophysiological recordings have been performed. **F**) Placement of a stainless-steel screw over the contralateral parietal bone, used as a reference during recordings.

**1.7.** Secure the upper incisors in the bite bar and gently pull the head backward to verify proper positioning. Reposition the incisors if any movement of the skull is detected.

**CRITICAL STEP:** Proper fixation of the head is essential for accurate stereotaxic targeting. Even slight skull movement may compromise injection accuracy.

**1.8.** Position the anesthesia nose cone and apply ophthalmic ointment to both eyes to prevent corneal drying.

**1.9.** Administer meloxicam (5-10 mg/kg, SC) for pre-emptive analgesia.

**1.10.** Position the ear bars symmetrically and apply 4% lidocaine gel to their tips before fixation.

**CRITICAL STEP:** Incorrect ear bar placement may cause skull misalignment and inaccurate stereotaxic coordinates.

**1.11.** Continuously monitor anesthetic depth throughout the procedure by assessing respiratory rate and the absence of pedal withdrawal reflexes.

**NOTE:** If spontaneous movements or reflexes are observed, increase the isoflurane concentration until surgical plane anesthesia is restored. If respiration ceases, immediately discontinue isoflurane administration and provide 100% oxygen until spontaneous breathing resumes.

### 2. Surgical field preparation and craniotomy

**NOTE:** Before starting the procedure, verify that the skull is firmly secured in the stereotaxic frame. All surgical procedures described below were performed under a surgical microscope with direct illumination.

**2.1** Remove the fur using depilatory cream, clean the skin with cotton swabs, and disinfect the surgical area with povidone-iodine (Fig. 2B).

**2.2** Apply 4% lidocaine gel to the incision site. Perform a midline skin incision using a scalpel (Fig. 2C) and extend it with fine scissors to fully expose bregma, lambda, the parietotemporal margins, and the occipital and interparietal bones (Fig. 2D). Carefully avoid damaging the underlying musculature.

**2.3** Clean and dry the skull to improve visualization of bregma and lambda.

**2.4** Using the glass micropipette prepared in step 1.2, identify bregma and lambda and level the skull in both the mediolateral and anteroposterior axes (Fig. 2D). Adjust the stereotaxic frame until the dorsoventral difference between landmarks is ≤ ±0.03 mm.

**CRITICAL STEP:** Accurate skull leveling is essential for reliable stereotaxic targeting. A dorsoventral deviation greater than ± 0.03 mm may significantly reduce targeting accuracy.

**2.5** Confirm skull alignment by measuring additional reference points located 1.5 mm posterior to bregma and ± 1.2 mm lateral. Readjust the ear bars, if necessary, until the dorsoventral difference remains ≤ ± 0.03 mm.

**2.6** Identify the desired stereotaxic coordinates according to the experimental design and perform the craniotomy using a surgical microdrill (Figure 2E). The following coordinates were used in this study: medial deep cerebellar nucleus (AP: −6.24 to -6.47 mm; ML: ±0.6–0.8 mm from bregma), cerebellar vermis (AP: −7.20 to −7.56 mm; ML: 0 mm from bregma), and cerebellar hemispheres (AP: −6.69 to −7.32 mm; ML: ±1.6 mm from bregma).

**CRITICAL STEP:** Hold the drill at approximately 45° relative to the skull surface and progressively thin the bone until it can be removed with fine forceps without damaging the dura mater.

**NOTE:** If bleeding occurs during the craniotomy, control it using sterile absorbent triangles. If hemostasis cannot be achieved, terminate the procedure and humanely euthanize the animal.

### 3. Stereotaxic viral delivery

**NOTE 1:** For all procedures, leave the micropipette in place for 5 min after injection before slowly withdrawing it to minimize viral reflux.

**NOTE 2**: The transduction efficiency of AAVs depends on many factors and remains difficult to predict, so an empirical approach is often required to determine the serotype and titer with the best performance in the desired cell population^37^.

#### 3.1 Pilot viral tracing procedures for CB projection mapping

The following pilot viral tracing experiments were performed in Ai14 mice to validate transduction of PCs and visualize PC projections to mDCN. Results were used to guide the location of viral injections for optogenetic experiments.

##### 3.1.1 Pilot 1: Local cerebellar cortical injections to label Purkinje cells

The goal of the first pilot experiment was to optimize injection coordinates for tagging PCs in CB vermis that project to mDCN. Ai14 mice, which conditionally express tdTomato, were injected locally with PHP.eB-hSyn-Cre-P2A-tdTomato in CB vermis, and tdTomato-labeled PC axonal projections in mDCN were confirmed histologically (Fig. 3**A1-5**).

**Figure 3:**
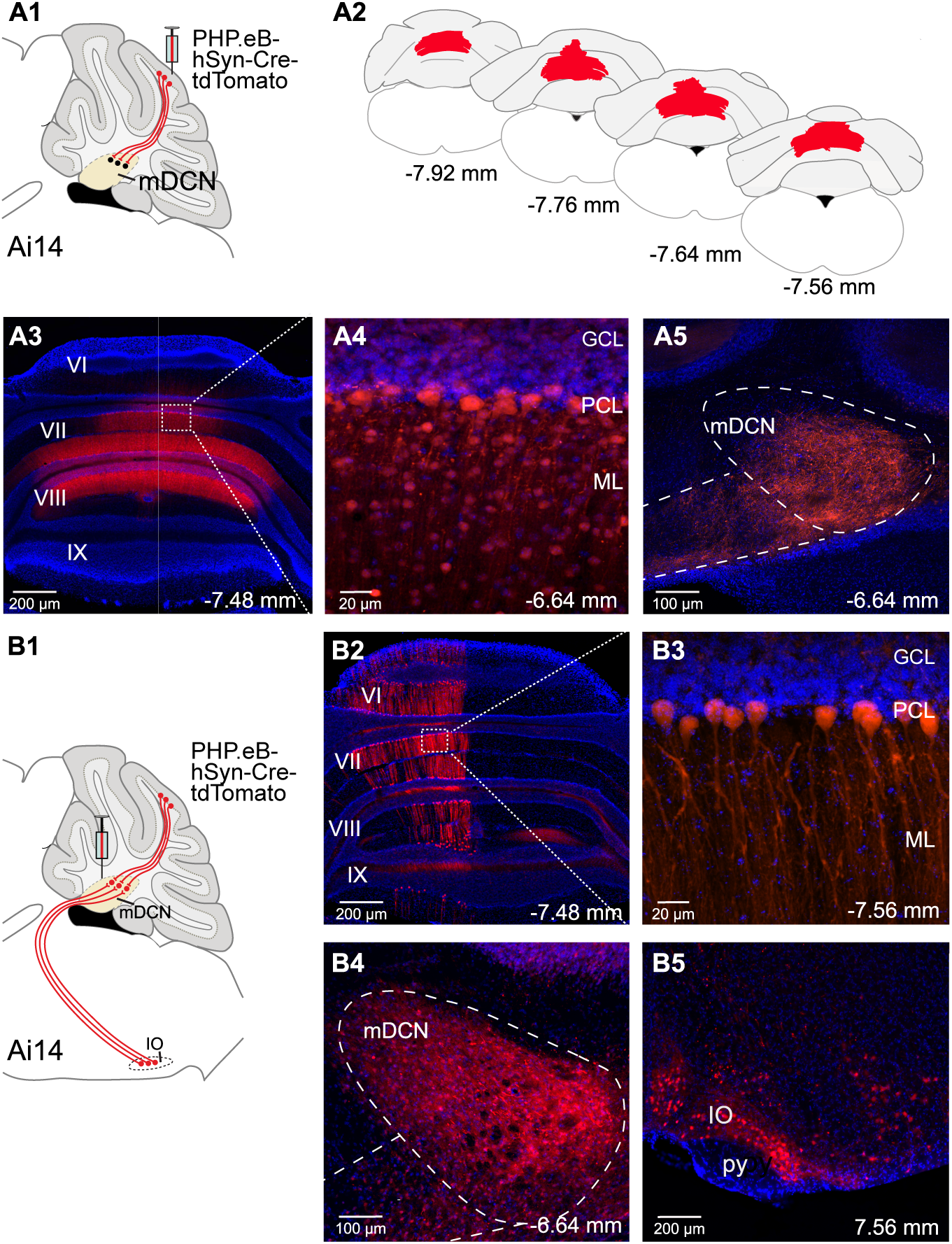
Anterograde and retrograde labeling approaches of PC-mDCN pathway. **A1)** Schematic of anterograde labeling approach. **A2**) Extent of viral spread in CB cortex, shown on consecutive coronal schematic sections from −7.92 to −7.56 mm relative to bregma. **A3**) Coronal section through the CB vermis showing labeled neurons. **A4**) Higher-magnification view of the boxed region in A3 with CB cortical layers annotated. **A5**) Labeled PC axons in mDCN. **B1**) Schematic of retrograde labeling approach. **B2**) Coronal section through CB vermis showing retrogradely labeled PCs. **B3**) Higher-magnification view of the boxed region in B2 with CB cortical layers annotated. **B4**) Coronal section through mDCN showing locally transduced neurons and labeled axons. **B5**) Retrogradely labeled neurons in the IO, which projects to mDCN. Blue: DAPI; Red: tdTomato. Abbreviations: GCL, granule cell layer; ML, molecular layer; PCL, Purkinje cell layer; mDCN, medial deep cerebellar nucleus; IO, inferior olive; py, pyramidal tract; VI–IX, lobules VI–IX of the CB vermis.

**3.1.1.a** After craniotomy at specified coordinates for vermis, back-fill a pulled micropipette with mineral oil, then tip-fill with virus (300 nL, see Table of Materials) using the stereotaxic microinjection system.

**3.1.1.b** Slowly lower the micropipette 1000 µm from brain surface and inject 0.45 µL of virus at 2 nL/s.

##### 3.1.2 Pilot 2: Combinatorial labeling of Purkinje cells from the medial deep cerebellar nucleus

The goal of the second pilot experiment was to optimize injection coordinates for retrograde labeling of PCs projecting to mDCN. Ai14 mice were injected in mDCN with PHP.eB-hSyn-Cre-P2A-tdTomato, and retrogradely tdTomato-expressing PCs in cerebellar vermis were identified histologically (Fig. 3**B1-4**).

**3.1.2.a** After craniotomy at the above specified coordinates for mDCN, back-fill a pulled glass micropipette with mineral oil, then tip-fill it with virus (300 nL) using the stereotaxic microinjection system.

**3.1.2.b** Slowly lower the micropipette 2,170 µm from brain surface and inject 250 nL of the virus at 2 nL/s.

**NOTE:** Because PHP.eB-Cre-P2A-tdTomato itself expresses tdTomato, its use in Ai14 mice does not permit independent validation of Cre activity. However, it remains suitable for optimizing injection coordinates for subsequent ChETA-EYFP experiments.

#### 3.2 Anterograde approach for opsin expression in Purkinje cell terminals

Following histological confirmation of coordinates for CB cortical regions projecting to mDCN (step 3.1.1), inject an AAV encoding the excitatory opsin ChETA (e.g., AAV-hSyn-hChR2(H134R)-EYFP) into the CB vermis at specified coordinates, at two depths (1000 µm and 600 µm from brain surface). (Fig. 4**A1**) Deliver 250 nL per injection site at 2 nL/s.

**Figure 4:**
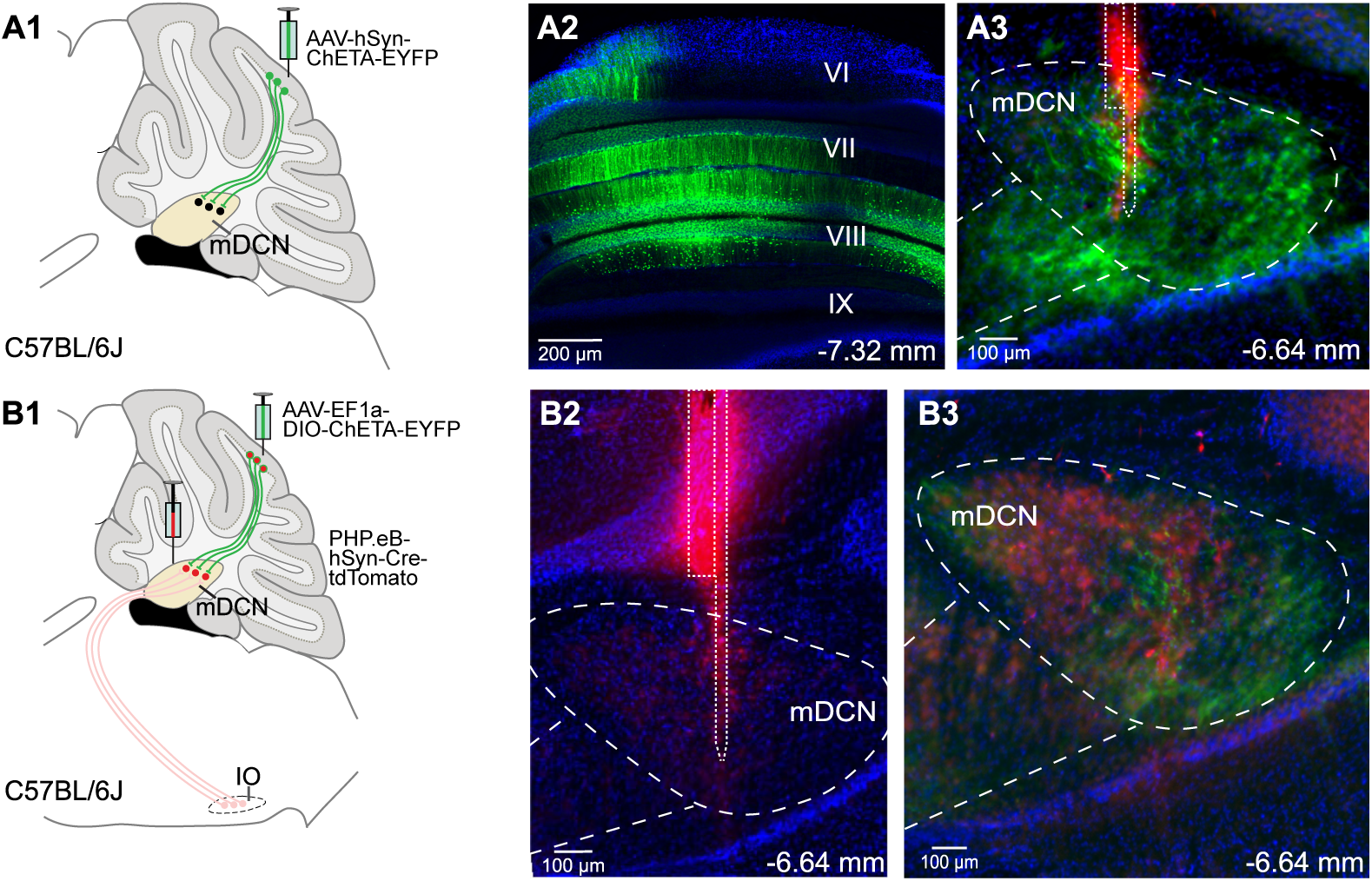
Viral expression of ChETA for optogenetic activation of PC inputs to mDCN. **A1**) Schematic of approach. **A2**) Coronal section of CB vermis showing ChETA-expressing neurons (green). **A3**) Coronal section through mDCN showing ChETA-EYFP expressing PC axons (green) and the location of the recording probe (red). **B1**) Schematic of approach. **B2**) Recording site in mDCN, with the probe track in red. **B3**) Coronal section through mDCN showing ChETA-EYFP-expressing PC axons (green) and AAV-PHP.eB-transduced neurons (red). Blue: DAPI. Abbreviations: mDCN, medial deep cerebellar nucleus; VI–IX, lobules VI–IX of CB vermis.

**NOTE**: General and neuron-specific promoters such as EF1a and hSyn, respectively, express well in CB but expression is not cell-type-specific^37,38^. However, because only PCs project out of CB cortex, optogenetic stimulation over mDCN selectively activates PC axon terminals.

#### 3.3 Combinatorial approach for projection-specific opsin expression in Purkinje cell terminals in medial deep cerebellar nucleus

To selectively express an excitatory opsin in PCs projecting to mDCN, inject a retrograde Cre-expressing viral vector (e.g., PHP.eB-hSyn-Cre-tdTomato) into mDCN and a Cre-dependent ChETA AAV (AAV-EF1a-DIO-hChR2(H134R)-EYFP-WPRE-HGH) into CB vermis (Fig. 4**B1-3**) (see Table of Materials). Perform all injections at 2 nL/s using the stereotaxic coordinates determined in step 3.1.

**NOTE**: In our hands, the widely used retrograde rAAV2^39,40^ does not infect PC terminals appreciably in C57BL/6J mice (not shown). Even though PHP.eB is typically delivered retro-orbitally for system-wide infection^41^, here we corroborate its suitability for local direct and/or retrograde transduction of PCs in C57BL/6 mice^42^, addressing the problem of retrograde viral tagging of PCs.

### 4. Postoperative care

**4.1** Upon completion of injection, slowly retract the injection pipette and proceed to close the incision with surgical sutures.

**4.2** Terminate anesthesia delivery and place the mouse on a 35 °C heating pad and continuously monitor until it fully regains consciousness and spontaneous locomotion. Inspect the sutured incision for bleeding before returning the mouse to the home cage.

**4.3** Administer buprenorphine (0.1 mg/kg, SC) immediately after surgery and meloxicam (5-10 mg/kg, SC) every 24 h during the first 2 postoperative days.

**4.4** Monitor animals twice daily during the first 48 h and at least once daily for 7 days thereafter. Assess body weight, wound healing, general condition, and signs of pain or distress.

**NOTE**: Administer additional buprenorphine (0.1 mg/kg, SC) every 8-12 h if signs of pain are observed. Animals exhibiting severe distress or body weight loss exceeding 20% should be humanely euthanized according to institutional animal care guidelines.

**4.5** Remove surgical sutures after complete wound healing, typically 7 days after surgery and no later than 14 days postoperatively.

### 5. Cranial access surgery for in vivo electrophysiological recordings

**NOTE**: Electrophysiological recordings were performed 4 weeks after viral vector injection. Unless otherwise indicated, all surgical procedures were identical to those described in Sections 1 and 2.

**5.1** Following induction of anesthesia with 2% isoflurane, administer an intraperitoneal anesthetic cocktail containing ketamine (100 mg/kg), xylazine (10 mg/kg), and acepromazine (1 mg/kg) before transferring the animal to the stereotaxic frame.

**5.2** Perform a craniotomy over mDCN using the stereotaxic coordinates for adult mice (AP: −6.45 mm; ML: ± 0.6 mm from bregma), following the procedure described in Section 2.

**5.3** Irrigate the craniotomy with sterile 0.9% saline, carefully remove the meninges using a 27 G hook-shaped needle, and rinse the exposed brain surface again with sterile saline.

**CRITICAL STEP**: Avoid damaging the brain surface while removing the meninges, as this may compromise recording quality.

**5.4** Perform a second craniotomy over the contralateral parietal bone and insert a stainless-steel screw (see Table of Materials) to serve as the reference electrode during electrophysiological recordings (Fig. 2F).

**CRITICAL STEP**: During the second craniotomy, hold the drill perpendicular (90°) to the skull surface. The appearance of a small swirl of bone dust around the drill bit indicates that the bone has been sufficiently thinned.

**NOTE**: If bleeding occurs, control it as described in Section 2.

### 6. Preparation for electrophysiological recordings and probe insertion

**NOTE:** Perform all procedures under a surgical microscope with direct illumination.

**6.1** Before discontinuing isoflurane administration, inject approximately 0.03 mL of the anesthetic cocktail described in Section 5.1. Repeat this dose every 30 min to maintain a stable anesthetic plane throughout the experiment.

**NOTE:** To minimize effects of ketamine and variable anesthetic depth on neuronal activity, administer each maintenance dose before the start of a recording and wait at least 10 min before data acquisition.

**6.2** Disconnect the animal from the isoflurane vaporizer, oxygen supply, and heating pad. Transfer the stereotaxic frame to a Faraday cage and reconnect the animal to a feedback-controlled heating pad and an oxygen supply (0.8–1.0 L/min). Confirm adequate anesthetic depth by assessing the absence of pedal withdrawal reflexes.

**6.3** Secure the reference wire to the stainless-steel screw described in Section 5.4 and insert the ground wire beneath the skin and musculature over the contralateral parietal bone.

**CRITICAL STEP:** Wrap the reference wire around the screw at least twice before securing it to ensure stable electrical contact and minimize environmental electrical noise.

**6.4** Mount the silicon probe onto the electrode holder, connect the Omnetics connector, and coat the probe tip with a fluorescent dye (see Table of Materials) to enable subsequent histological verification of the recording site.

**6.5** Position the probe above the craniotomy and connect the acquisition system, stimulation controller, and laser (see Table of Materials). Start the acquisition software to monitor the electrophysiological signal in real time.

**6.6** Slowly lower the probe until it contacts the brain surface.

**NOTE:** Contact with the brain surface can be identified by the abrupt transition from background noise to a stable local field potential (LFP) signal.

**6.7** After advancing the probe approximately 100 µm into the CB, switch off the illumination to minimize 60 Hz electrical noise and continue lowering the probe manually at approximately 1 µm/s until the desired recording depth is reached (approximate DV = 3.0 mm for mDCN).

**CRITICAL STEP:** Avoid insertion speeds greater than 1 µm/s, as rapid probe advancement may cause tissue damage, reduce recording quality, and compromise subsequent spike sorting.

### 7. Example optogenetic stimulation and recording protocol

**7.1** After the probe reaches the target recording depth, allow the preparation to stabilize for at least 20 min before starting the recordings.

**NOTE**: Confirm correct probe placement by monitoring neuronal activity. Under ketamine anesthesia, neurons in mDCN typically exhibit alternating periods of high firing activity and silence^43^ (Fig. 5A), in contrast to the more continuous activity recorded in the cerebellar cortex^44^ (Fig. 5B).

**Figure 5:**
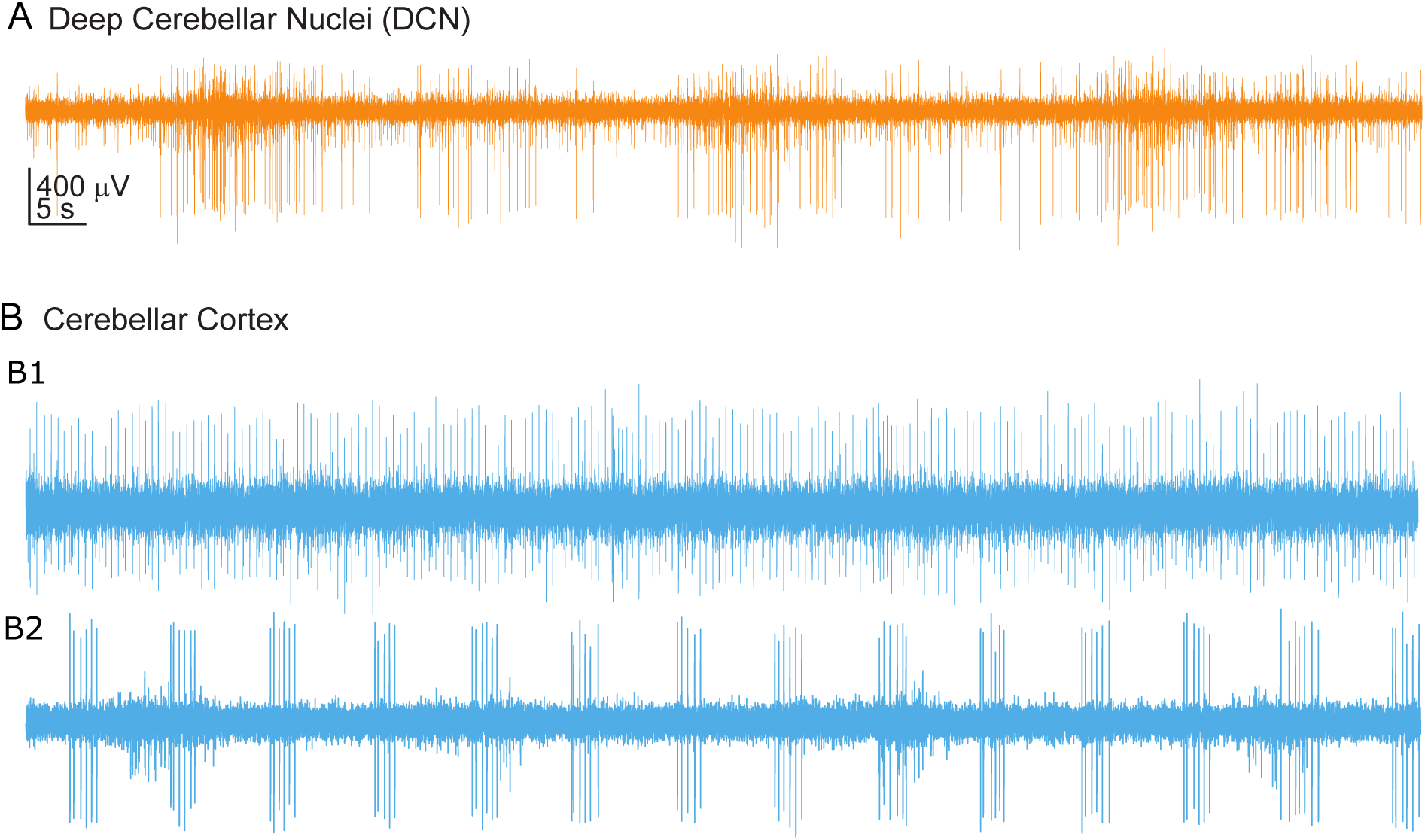
Comparison of electrophysiological signals recorded from deep cerebellar nuclei and cerebellar cortex of anesthetized mouse. **A**) Representative electrophysiological trace recorded from DCN, showing an irregular firing pattern characterized by periods of increased neuronal activity and/or bursting, interspersed with periods of reduced or absent activity. **B1-2**) Representative electrophysiological traces recorded from CB cortex, showing more regular firing patterns compared to DCN.

**7.2** Configure the optogenetic stimulation protocol using the acquisition software to control the pulse generator, which delivers TTL pulses to trigger the laser.

**NOTE 1**: Administer ketamine maintenance doses immediately before each recording session and allow at least 10 min before data acquisition to minimize ketamine-induced electrophysiological effects^45,46^.

**NOTE 2**: Use a laser pulse duration of 3 ms, as this is sufficient to reliably evoke a single action potential in single PCs during optogenetic stimulation ex vivo^47^

**7.3** Apply stimulation trains at 1, 10, and 20 Hz, each delivered at 0.1, 1, and 10 mW laser power (measured at end of optic fiber). Configure the pulse generator according to Table 1 **[Place Table 1 here].**

**7.4** Record a 90-s baseline period followed by 20 s of recording, consisting of 1 s of stimulation and 19 s of inter-burst interval. Repeat the protocol for 10 trials at each laser intensity, maintaining constant inter-recording intervals.

**NOTE**: Acquire all three laser intensities during the same recording session to avoid performing spike sorting separately for each stimulation condition.

### 8. Post-recording tissue collection and processing

**8.1** At the end of the recording session, euthanize the animal with an overdose of pentobarbital (≥100 mg/kg, IP; see Table of Materials; **CAUTION**: Pentobarbital is a controlled euthanasia agent. Handle and dispose of it according to institutional and governmental regulations) while maintaining deep anesthesia. Confirm the absence of reflexes before proceeding

**8.2** Perform transcardial perfusion with approximately 10 mL of phosphate-buffered saline (PBS), followed immediately by 4% paraformaldehyde (PFA) in PBS (**CAUTION**: PFA is toxic and a potential irritant. Prepare and handle all PFA solutions in a certified chemical fume hood while wearing appropriate personal protective equipment). Remove the brain and postfix it in 4% PFA for 24 h at 4 °C.

**8.3** Dissect the cerebellum, mount onto a vibratome stage using cyanoacrylate adhesive (**CAUTION**: Cyanoacrylate adhesive bonds skin and eyes rapidly. Avoid direct contact) and cut 80 µm coronal sections.

**8.4** Store sections in PBS containing 0.01% sodium azide (**CAUTION**: Sodium azide is highly toxic and may form explosive metal azides in plumbing. Dispose of sodium azide-containing solutions according to institutional hazardous waste regulations) until histological processing.

**8.5** Mount sections onto microscope slides using a DAPI-containing mounting medium (see Table of Materials), apply top coverslips, and seal the edges with clear nail polish.

### 9. Histological verification of viral expression and probe placement

**9.1** Image the processed brain sections using an epifluorescence microscope (e.g., Keyence BZ-X1000; see Table of Materials) to verify viral expression and recording probe placement.

**9.2** Identify and map the anatomical structures and recording sites according to the mouse brain atlas by Paxinos and Franklin.

### 10. Electrophysiological data analysis

#### 10.1 Spike sorting

**NOTE 1**: Perform spike sorting using Kilosort4^48^ followed by manual curation in Phy2^49^. Our protocol was implemented under Windows 10 using Open Ephys^50^ *continuous* files, and equivalent procedures can be applied to other operating systems and file formats.

**10.1.1** Create and activate a Kilosort4 Anaconda environment, then launch the graphical interface:

conda activate kilosort python -m kilosort

**10.1.2** Convert the Open Ephys *.continuous* files to a binary (*.bin*) file using the **Convert to Binary** tool. Select **Open Ephys** as the file type, **CH** as both *stream_id* and *stream_name*, and **int16** as the data type.

**CRITICAL STEP:** Manually append the .bin extension to the output filename before conversion. Otherwise, the file will not be recognized by Kilosort.

**10.1.3** Load the probe configuration. If the probe is not included in the default library, generate a custom probe layout following the Kilosort documentation. The probe configuration used in this study is provided in **Supplementary File X**.

**10.1.4** Load the binary recording, specify the output directory, and configure the acquisition parameters (32 channels, 30 kHz, int16).

**CRITICAL STEP:** Verify the probe-dependent parameters (*dmin*, *dminx*, and *min_template_size*) before running the algorithm, as these values strongly influence spike sorting performance.

**10.1.5** Run Kilosort4. Upon completion, a kilosort4 folder containing all files required for manual curation will be generated.

**10.1.6** Manual curation of single units

**NOTE:** Detailed documentation for Phy2 is available at the official documentation website and a step-by-step video tutorial describing the manual curation workflow can be found at: https://www.youtube.com/watch?v=czdwIr-v5Yc

**10.1.7** Open **Anaconda Prompt**, activate the Phy2 environment, and set the working directory to the kilosort4 output folder:

conda activate <ENVIRONMENT_NAME>

cd <PATH_TO_KILOSORT4>

phy template-gui params.py

**10.1.8** Inspect each cluster by evaluating waveform shape, auto- and cross-correlograms, refractory period violations, spatial distribution across recording channels, and temporal stability.

**10.1.9** Merge or split clusters when necessary, discard noise or poorly isolated units, and label well-isolated single units as “good” for subsequent analyses.

#### 10.2 Offline data analysis in MATLAB

**10.2.1** Import the .*continuous* electrophysiological files into MATLAB using the Open Ephys Load Faster toolbox. Load the recordings, timestamps, stimulation channel, and spike-sorting output generated by Kilosort4 and manually curated with Phy2.

**10.2.2** Preprocess the electrophysiological signals by applying notch filters at 50 Hz or 60 Hz, depending on the local power-line frequency, to reduce electrical interference. Apply additional filtering steps as needed to improve signal quality.

**10.2.3** Detect the onset of each optogenetic stimulation train from the analog input channel used to record the laser TTL signal. Use these timestamps to align neuronal activity to the onset of each stimulus train.

**NOTE:** Detect stimulation onset from the recorded TTL signal rather than from the programmed stimulation protocol to account for potential timing offsets between the stimulation and acquisition systems.

#### 10.3 Quantification of evoked responses

Laser pulses were classified according to their order within each stimulus train (1st, 2nd, 3rd, etc.), and trials were grouped by stimulation condition and pulse order. For each pulse, three epochs were defined: baseline (30 ms before pulse onset), pulse (3 ms), and rebound (27 ms after pulse offset).

Firing rate was calculated as:

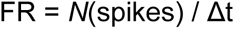

where N(spikes) is number of spikes observed during every single bin, and Δt is the 0.0005-s bin size. Firing rate was therefore in units of spikes/second (Hz). The same procedure was followed to quantify firing rate during the baseline and rebound epochs.

The percentage change in firing rate relative to baseline (ΔFR) was calculated as:

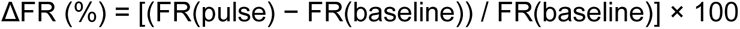

Values were averaged across trials for each pulse. The rebound window was calculated following each individual pulse rather than after the end of the stimulation train.

#### 10.4 Construction of the PSTH

For each pulse, a peri-stimulus time histogram (PSTH) was generated using 0.5-ms bins within a ±30-ms window centered on pulse onset. Firing rate for each bin was calculated as:

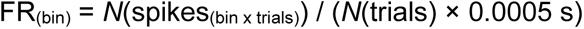

Values were expressed directly in Hz without additional normalization or smoothing.

#### 10.5 Change in firing probability

For each 0.003-s stimulation pulse, the number of 0.0005-s bins that had at least one spike was counted and added to all other pulses in a given trial, and all other trials in a given neuron and stimulation intensity. Therefore, the observed firing probability was calculated as:

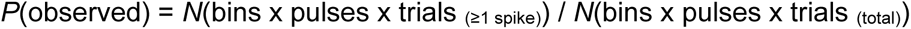

Baseline firing probability was computed the same way during the 0.030-s period preceding each pulse:

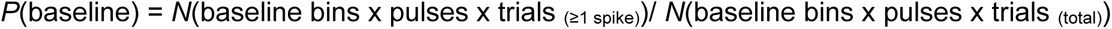

Firing probability during the stimulation pulses was normalized to the baseline firing rate:

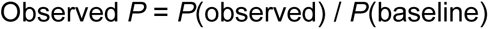

No difference from the pre-pulse baseline firing probability would be expected to result in Observed *P* = 1. Values of Observed *P* > 1 indicate an increase in firing probability relative to pre-pulse baseline, whereas values < 1 indicate a decrease. Employing the same bin size during the baseline as during the stimulation pulse ensures unbiased analysis of the two windows.

#### 10.6 Statistical analysis

**10.6.1** Export the processed data from MATLAB and import the resulting files into RStudio for statistical analysis. Compare firing rate across baseline, stimulation, and rebound periods using Friedman tests followed by Wilcoxon signed-rank post hoc tests with Holm correction.

**10.6.2** Analyze the effects of laser intensity (0.1, 1, and 10 mW) and stimulation frequency (1, 10, and 20 Hz) on ΔFR and relative firing probability using separate 3 × 3 permutation-based factorial ANOVAs with 10,000 permutations and the Freedman–Lane procedure. Perform post hoc comparisons using independent-samples Wilcoxon tests with Holm correction. For relative firing probability, additionally compare each condition with the baseline reference value of 1 using one-sample Wilcoxon tests with Holm correction.

**10.6.3** Consider results statistically significant at *p* < 0.05.

**Note**: All analysis scripts, processed data files, and the underlying electrophysiological recording data used for these analyses will be made available in the repository at http://github.com/FioravanteLab/2026_Merino_JoVE to enable replication of the analysis pipeline and results.

## REPRESENTATIVE RESULTS

### Pilot viral tracing experiments and optimization of stereotaxic coordinates

Before conducting the optogenetic experiments, two pilot tracing experiments were performed to optimize injection coordinates and confirm anterograde and retrograde viral infection of PCs. In the first pilot experiment, Ai14 mice (N = 4 females) were injected with a PHP.eB-hSyn-Cre viral vector (1.9 x 10^13 gc/ml) locally into the CB vermis at specified coordinates (Figs. 3**A1-2**). Twenty-one days later, abundant tdTomato+ neurons were detected in CB cortical slices (Fig. 3**A3**), including PCs identified based on cell body size and location in Purkinje cell layer (Fig. 3**A4**). tdTomato+ PC axons could be readily visualized in mDCN (**Fig. 3A5**). These results confirm cortical injection coordinates and document the ability of the PHP.eB serotype to transduce local PCs in C57BL/6J mice. We have previously confirmed the ability of other AAV serotypes (not shown; and ref.^37,51^) to transduce PCs.

In the second pilot experiment, Ai14 mice (N = 2 females) were injected unilaterally with PHP.eB-hSyn-Cre-P2A-tdTomato in mDCN at specified coordinates (Fig. 3**B1**). At a titer of 1.9 x 10^13 gc.ml, retrogradely labeled PCs were readily identified mainly in posterior vermis, between −7.56 and −7.92 mm from bregma (Figs. 3**B2-3**), and tdTomato+ axons were evident in mDCN (Fig. 3**B4**). Retrogradely labelled neurons were also detected in inferior olive (Fig. 3**B5**), which provides climbing fiber innervation to DCN^52^. Diluting the titer 10-fold also results in effective retrograde PC transduction while minimizing retrograde spread to inferior olive (not shown). These results confirm mDCN injection coordinates and validate the ability of PHP.eB to retrogradely transduce PCs (and inferior olive neurons at high titers) in these mice. The anatomical information obtained from these pilots defined the stereotaxic coordinates for the viral injections and electrophysiological recordings reported below.

### Optogenetic stimulation of Purkinje cell axon terminals in medial deep cerebellar nuclei

**Anterograde approach.** We monitored the firing rate of mDCN neurons in response to optogenetic stimulation of axonal terminals of PCs anterogradely labeled with ChETA, using stimulus trains of varying frequencies (1, 10 and 20 Hz) and laser light intensities (0.1, 1 and 10 mW). All statistical values for these experiments are provided in Supplementary Table 1. At 0.1 mW, we did not observe significant firing rate modulation at any stimulation frequency (Friedman test; 1, 10, and 20 Hz: *p* = 0.744, 0.249, and 0.216, *n* = 44, 44, and 32 units, respectively; *N* = 3 mice, both sexes) (Fig. 6A-B). In contrast, stimulation with 1 and/or 10 mW light produced robust firing rate suppression at all frequencies tested (1 mW light, 1, 10, and 20 Hz: all *p* < 0.0001, *n* = 45 units; 10 mW light, 1, 10, and 20 Hz: all *p* < 0.0001, *n* = 57). Post-hoc pairwise comparisons using the Wilcoxon signed-rank test with Holm correction confirmed a significant reduction in firing rate during the light pulse relative to baseline across all six intensity-frequency conditions (all *p* < 0.0001). Following cessation of stimulation, firing rate increased significantly relative to the stimulation period (Pulse vs. Rebound, all *p* < 0.0001), indicating recovery toward baseline values. Comparisons between baseline and rebound periods revealed no significant differences in any condition (1 mW, 1, 10, and 20 Hz: *p* = 0.797, 0.746, and 0.780, respectively; 10 mW, 1, 10, and 20 Hz: *p* = 0.987, 0.280, and 0.867, respectively) (Figs. 6A-B).

**Figure 6:**
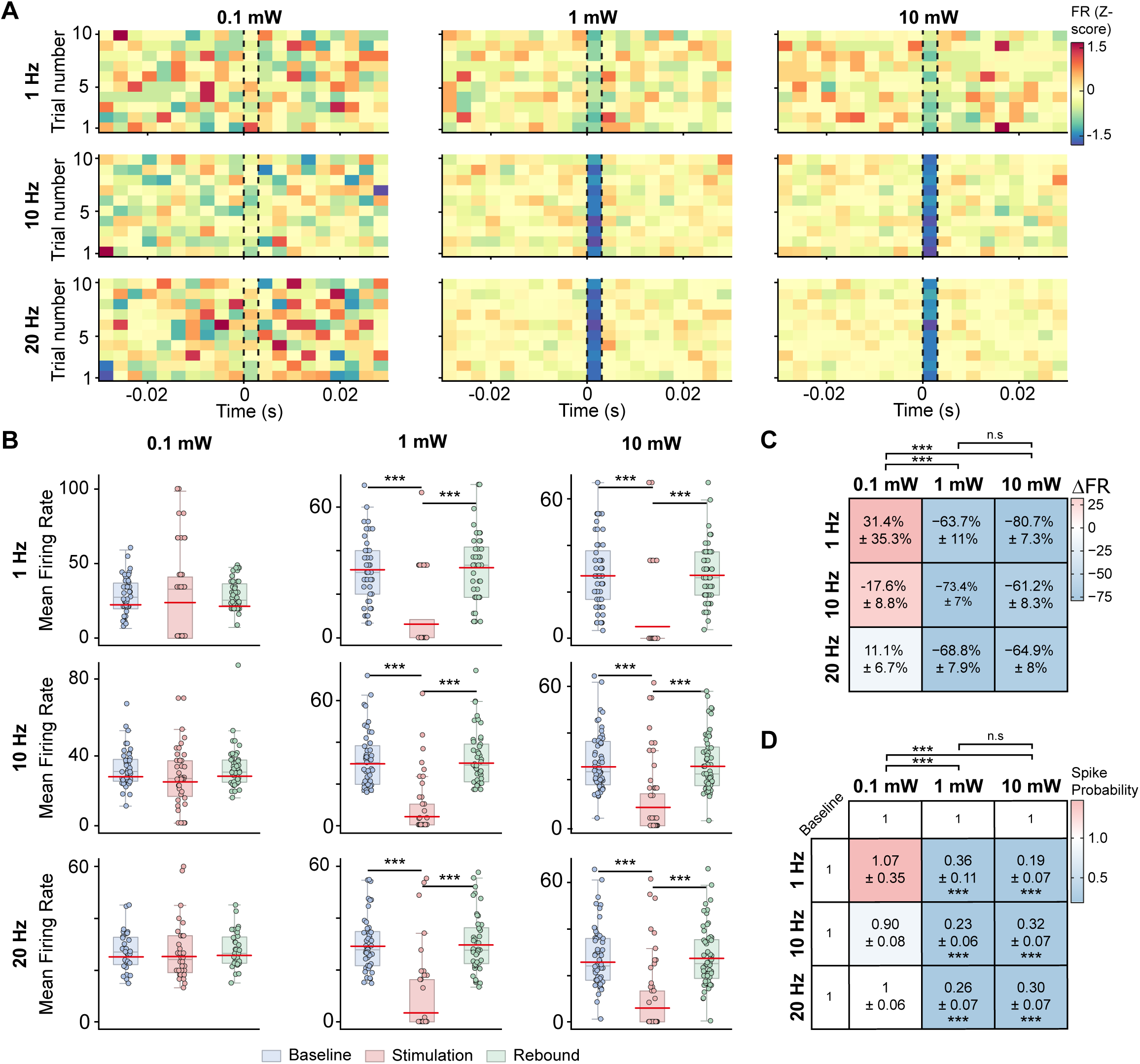
Electrophysiological responses of medial DCN neurons to optogenetic stimulation of Purkinje cell axon terminals labeled with anterograde approach. A) Average normalized (z-score) firing rate heatmaps showing neuronal responses to optogenetic stimulation across three stimulation frequencies (1, 10, and 20 Hz) and three laser intensities (0.1, 1, and 10 mW). Each heatmap contains 10 rows corresponding to the 10 stimulation trials performed for each frequency × intensity condition. For each trial, the heatmap shows a 30-ms baseline period, a 3-ms stimulation period, and 30 ms following stimulation (the rebound period). For 10- and 20-Hz stimulation, responses to individual 3-ms laser light pulses delivered during the 1-s stimulation period were averaged. B) Quantification of firing rate during baseline, stimulation, and rebound periods for each frequency × intensity condition. Box plots show the median, first and third quartiles, and interquartile range; C) Mean percent change in firing rate (ΔFR) between baseline and stimulation for each frequency × intensity condition. D) Relative firing probability during stimulation compared with baseline for each frequency × intensity condition. Values of 1 represent the baseline reference. Asterisks within the heatmap indicate conditions significantly different from baseline (one-sample Wilcoxon test against 1, Holm-corrected). * p < 0.05; ** p < 0.01; *** p < 0.001.

To further evaluate the effects of laser intensity and stimulation frequency on neuronal responses, factorial analyses were performed separately for percent change in firing rate (ΔFR; Fig. 6C) and relative firing probability (Fig. 6D). A permutation-based 3 × 3 factorial ANOVA (Freedman-Lane procedure, 10,000 permutations) was used in both cases, as the data were not normally distributed and contained extreme values. Post-hoc comparisons were performed using independent-samples Wilcoxon tests with Holm correction. ΔFR relatively to baseline was analyzed with a 3 × 3 permutation-based factorial ANOVA. Laser intensity had a significant main effect (*p* < 0.001), indicating that the magnitude of the change in firing rate depended on stimulation intensity. The main effect of stimulation frequency did not reach significance (*p* = 0.054), but a significant frequency × intensity interaction was detected (*p* = 0.001), indicating that the influence of frequency on ΔFR differed across intensity levels. Post-hoc comparisons on the intensity main effect showed significant differences between 0.1 and 1 mW (*p* < 0.001) and between 0.1 and 10 mW (*p* < 0.001), whereas no significant difference was observed between 1 and 10 mW (*p* = 0.642). In summary, increasing laser intensity from 0.1 to 1.0 mW produced a marked reduction in DCN firing rate, with no further change between 1 and 10 mW, whereas stimulation frequency did not have a consistent effect on its own (Fig. 6C).

Relative firing probability (Fig. 6D) was analyzed as a function of stimulation intensity and frequency using the same permutation-based factorial ANOVA. Laser intensity had a significant main effect (*p* < 0.001), and significant effects of stimulation frequency (*p* = 0.023) and of the frequency × intensity interaction (*p* = 0.003) were also detected. Post-hoc comparisons on the intensity main effect showed significant differences between 0.1 and 1 mW (*p* < 0.001) and between 0.1 and 10 mW (p < 0.001), whereas no significant difference was observed between 1 and 10 mW (*p* = 0.632). In contrast, none of the pairwise comparisons between frequencies survived correction (1 vs. 10 Hz: p = 0.262; 1 vs. 20 Hz: *p* = 0.428; 10 vs. 20 Hz: *p* = 0.777).

To determine which stimulation conditions differed from baseline, each of the nine *frequency × intensity* combinations was additionally compared with the reference value of 1 using one-sample Wilcoxon tests with Holm correction. Relative firing probability was significantly reduced at 1 mW for 1, 10, and 20 Hz (p < 0.0001 for all comparisons) and at 10 mW for 1, 10, and 20 Hz (*p* < 0.0001 for all comparisons). None of the conditions at 0.1 mW differed significantly from baseline (1, 10, and 20 Hz: *p* = 1.000, 0.579, and 1.000, respectively). These results indicate that laser stimulation reduced firing DCN probability at 1 and 10 mW, whereas stimulation frequency had, at most, a modest, intensity-dependent modulatory effect (Fig. 6D).

Together, the firing rate and firing probability analyses indicate that optogenetic activation of PC axon terminals produces a robust, laser light intensity-dependent suppression of neuronal activity in mDCN. Stimulation frequency contributed only modestly in a manner that depended on laser intensity.

### Combinatorial approach

We expressed ChETA selectively in the subset of PCs that project to mDCN using a combinatorial approach, which leveraged retrograde infection with PHP.eB-Cre in mDCN and Cre-dependent (floxed) expression of ChETA in CB vermis, and monitored the firing rate of mDCN neurons in response to optogenetic stimulation of axonal terminals of PCs in mDCN. All statistical values for these experiments are provided in Supplementary Table 2. Similarly to results obtained with the anterograde approach, optogenetic stimulation did not significantly alter neuronal firing rate at 0.1 mW at any of the frequencies tested (Friedman test; 1, 10, and 20 Hz: *p* = 0.475, 0.145, and 0.368, respectively, *n* = 57 units; *N* = 3 mice, both sexes). In contrast, stimulation at 1 and 10 mW produced a significant and reproducible suppression of firing rate during the light pulse at all three frequencies tested (Friedman test; 1 mW light, 1, 10, and 20 Hz: all *p* < 0.0001, n = 51 units; 10 mW light, 1, 10, and 20 Hz: all *p* < 0.0001, n = 52 units). Post-hoc Wilcoxon signed-rank tests with Holm correction confirmed a significant reduction in firing rate during the light pulse relative to baseline in all six intensity-frequency conditions (all *p* < 0.001). Following cessation of the light stimulus, firing rate significantly increased relative to the stimulation period (Pulse vs. Rebound, all *p* < 0.001), consistent with recovery toward baseline values. Comparisons between baseline and rebound periods revealed no significant differences in most conditions (1 mW: *p* = 0.292, 0.907, and 0.725 for 1, 10, and 20 Hz, respectively; 10 mW: *p* = 0.725 and 0.859 for 1 and 10 Hz, respectively), although a small but significant baseline-rebound difference was observed with the strongest stimulation regime (10 mW-20 Hz; *p* = 0.039) (Figs. 7A-B).

**Figure 7:**
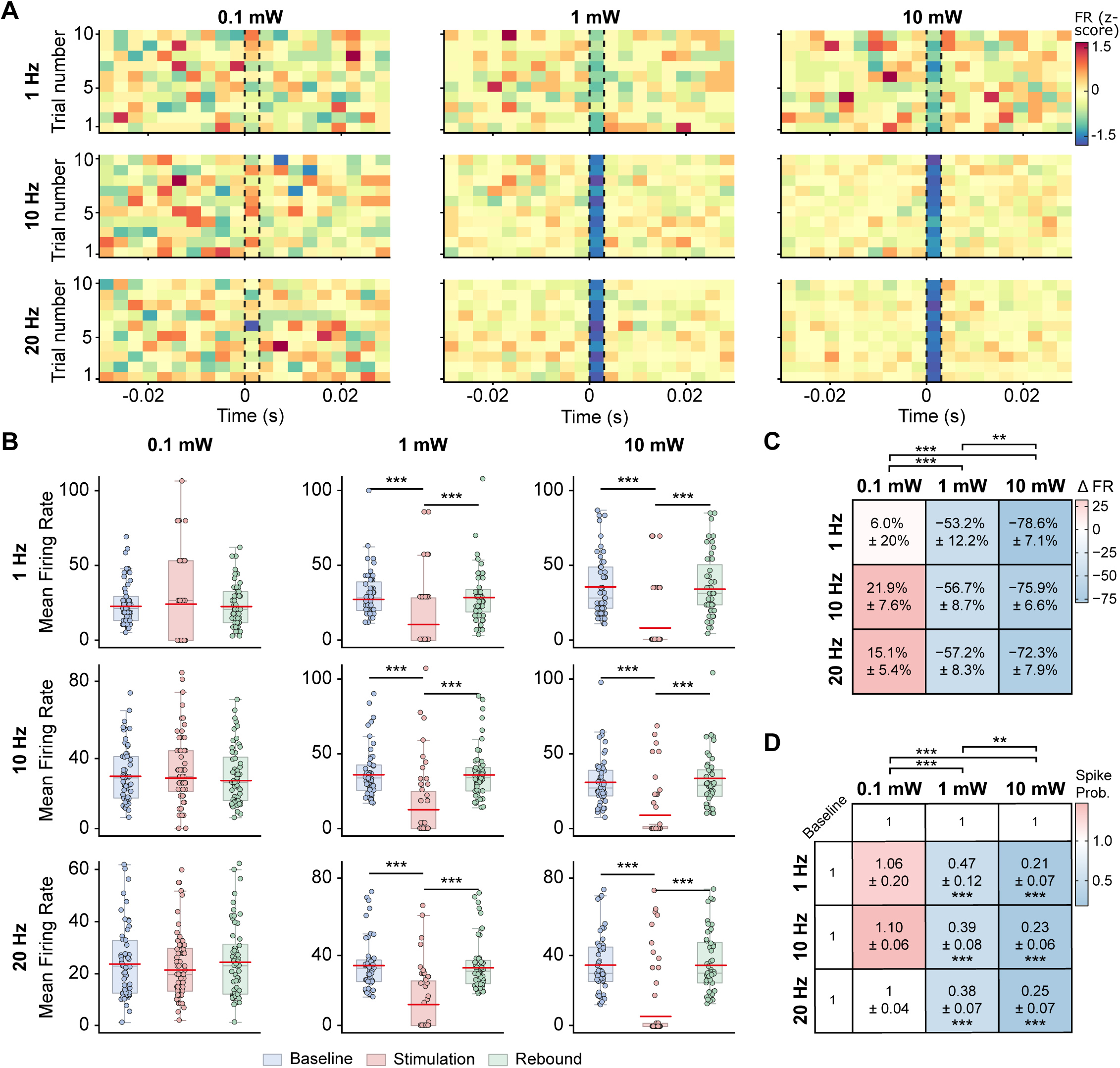
Electrophysiological responses of medial DCN neurons to Purkinje cell axon terminals labeled with combinatorial approach. **A**) Average normalized firing rate heatmaps showing neuronal responses to optogenetic stimulation across three stimulation frequencies (1, 10, and 20 Hz) and three laser intensities (0.1, 1, and 10 mW). Each heatmap contains 10 rows corresponding to the 10 stimulation trials performed for each frequency × intensity condition. For each trial, the heatmap shows a 30-ms baseline period, a 3-ms stimulation period, and the rebound period following stimulation. For 10- and 20-Hz stimulation, responses to individual 3-ms laser light pulses delivered during the 1-s stimulation period were averaged. **B**) Quantification of firing rate during baseline, stimulation, and rebound periods for each frequency × intensity condition. Box plots show the median, first and third quartiles, and interquartile range; red lines indicate the mean. **C**) Mean percent change in firing rate (ΔFR) between baseline and stimulation for each frequency × intensity condition. **D**) Relative firing probability during stimulation compared with baseline for each frequency × intensity condition. Values of 1 represent the baseline reference. Asterisks within the heatmap indicate conditions significantly different from baseline (one-sample Wilcoxon test against 1, Holm-corrected). * p < 0.05; ** p < 0.01; *** p < 0.001.

Factorial analyses were performed to evaluate the effects of laser intensity and stimulation frequency on ΔFR and relative firing probability, using a permutation-based 3 × 3 factorial ANOVA (Freedman–Lane procedure, 10,000 permutations) followed by independent-samples Wilcoxon post-hoc comparisons with Holm correction. No significant main effect of stimulation frequency was observed on ΔFR (*p* = 0.827), and there was no significant frequency × intensity interaction (*p* = 0.899). In contrast, laser intensity had a significant main effect (*p* < 0.001). Post-hoc comparisons showed significant differences between all three intensities: 0.1 vs. 1 mW (*p* < 0.001), 0.1 vs. 10 mW (*p* < 0.001), and 1 vs. 10 mW (*p* < 0.01). Thus, laser intensity significantly affected ΔFR at all three levels, whereas stimulation frequency did not significantly modify this effect (Fig. 7C).

Relative firing probability was similarly analyzed as a function of stimulation intensity and frequency. No significant main effect of stimulation frequency was observed (*p* = 0.870), and there was no significant *frequency × intensity* interaction (*p* = 0.943). In contrast, laser intensity had a significant main effect (*p* < 0.001). Post-hoc comparisons showed significant differences between 0.1 and 1 mW (*p* < 0.001), 0.1 and 10 mW (*p* < 0.001), and 1 and 10 mW (*p* < 0.01). Comparisons with the reference value of 1 further showed a significant reduction in relative firing probability under all conditions at 1 mW (1 Hz: *p* < 0.001; 10 and 20 Hz: *p* < 0.0001) and at 10 mW (1, 10, and 20 Hz: *p* < 0.0001 for all comparisons), whereas none of the conditions at 0.1 mW differed significantly from baseline (1, 10, and 20 Hz: *p* = 0.714, 0.553, and 0.714, respectively) (Fig. 7D).

Together, these results show that selective optogenetic activation of PC terminals projecting to mDCN produces a robust suppression of neuronal activity that depends primarily on laser intensity, with no detectable effect of stimulation frequency.

## DISCUSSION

This protocol combines two complementary AAV-based strategies for targeting PC-mDCN projections with localized optogenetic stimulation and simultaneous in vivo extracellular electrophysiological recordings. Both strategies produced robust suppression of mDCN neuronal firing during optogenetic activation of PC axonal terminals, consistent with the established inhibitory influence of PCs on CB nuclear neurons. These results demonstrate that both viral approaches enable effective manipulation of PC-mediated inhibition and provide a functional readout of its impact on CB output. Although both strategies produced similar physiological effects under the conditions tested, they differ in their targeting principles, experimental requirements, and the anatomical specificity they provide.

### Two viral strategies for targeting Purkinje cells: advantages and limitations

The first strategy employs localized AAV delivery to the CB cortex without requiring a cell-type-specific promoter or genetic driver. Although opsin expression is not restricted to PCs, selective stimulation of PC axonal terminals within mDCN can be achieved by exploiting the anatomical organization of CB cortical output. This approach offers experimental simplicity and flexibility in selecting the cortical region of origin, such as the vermis, CB hemispheres, or even specific lobules, while restricting optical stimulation to axonal projections within the targeted nuclear territory. In contrast, the combinatorial strategy restricts opsin expression to a specific PC subset defined by its mDCN projection and anatomical position in CB cortex, providing additional specificity. This projection-defined targeting requires two stereotaxic viral injections and accurate targeting of both the CB cortical region and the downstream nucleus, increasing experimental complexity. Importantly, neither strategy requires transgenic mouse lines, avoiding the expense and time associated with colony establishment and breeding while facilitating adaptation to other animal species. Thus, the choice between these approaches depends on whether the experimental objective requires selective stimulation of PC terminals within a defined nuclear territory or restriction of opsin expression to a specific PC projection.

Previous studies have demonstrated the utility of PC terminal photostimulation in both anesthetized preparations and awake, freely moving animals, enabling investigations of CB nuclear physiology and motor behavior^32–35^. The viral strategies described here can be adapted to these different experimental conditions and extended to other CB regions and animal models. Some important limitations should be considered when adapting the protocol: viral transduction efficiency and tropism can vary across animal strains and species^53–55^, while incomplete transduction may limit the proportion of the targeted PC population recruited during optical stimulation. In particular, the enhanced transduction efficiency of PHP.eB following systemic administration depends on the LY6A receptor and varies across mouse strains^56,57^. Although this dependence should not be directly extrapolated to local viral delivery, transduction efficiency should be validated when adapting the protocol to other strains or species^58^. Furthermore, retrograde viral delivery to the DCN can result in transduction of additional afferent populations, including neurons in the inferior olive (Fig. 3**B5**). The combinatorial strategy addresses this limitation by restricting delivery of the Cre-dependent opsin to the CB cortex, thereby limiting ChETA expression to neurons within the cortical injection site that are also transduced by the retrograde vector. However, neither approach distinguishes molecularly or physiologically distinct PC populations within the targeted CB cortical region or projection pathway. These considerations should guide the selection and validation of viral targeting strategies when adapting the protocol to other CB circuits or animal models.

### Critical experimental steps

Successful implementation of this protocol depends on accurate viral targeting, sufficient opsin expression at PC axonal terminals, and precise positioning of the optrode within mDCN. Stereotaxic accuracy is particularly important given the small size and deep location of mDCN. Careful leveling of the skull, empirical adjustment of injection and recording coordinates, and post hoc histological verification are therefore essential for reliable targeting. Proper execution of the surgical and anesthetic procedures is also essential for reproducibility. The injection volume and rate used here were selected to limit unwanted viral spread and tissue damage while achieving sufficient opsin expression. During electrophysiological recordings, ketamine/xylazine provides sustained anesthesia for prolonged procedures. Repeated maintenance doses allow anesthetic depth to be maintained throughout the recording session, and the 10-min wait period after each ketamine maintenance dose helps minimize confounding acute changes in LFP and single-unit activity associated with ketamine administration^45,59,60^. Additionally, slow probe advancement and sufficient stabilization time minimize tissue displacement and recording drift. Optical stimulation parameters must also be optimized to reliably suppress ongoing mDCN firing while minimizing post-inhibitory rebound firing. In the present protocol, we use brief light pulses (3 ms) and find that stimulation intensities of 1 and 10 mW, but not 0.1 mW, reliably suppress mDCN neuronal firing (Figs. 6,7). Together, these considerations are critical for obtaining reproducible electrophysiological responses and distinguishing successful anatomical targeting and viral expression from effective functional manipulation of the PC-mDCN circuit.

### Significance and potential applications

The ability to selectively manipulate anatomically defined PC populations and characterize their influence on downstream neural activity provides opportunities to investigate the contribution of CB cortical output to motor and nonmotor functions. This approach can be adapted to target PC projections to different DCN nuclei and combined with recordings in awake behaving animals, to determine how PC-mediated inhibition shapes CB output during motor learning, reward processing, prediction updating, and other behaviors. It may also be useful for investigating alterations in PC-DCN communication associated with neurological and neurodevelopmental disorders. For example, distinct patterns of DCN activity have been associated with ataxia, dystonia, and tremor, and experimentally reproducing specific patterns of altered SCN activity can induce related motor phenotypes in otherwise healthy animals^33,35^. DCN output also contributes to reward processing, fear learning, and social behavior^61–65^. The present protocol provides a means to investigate how anatomically defined PC populations influence the activity of cerebellar output neurons, establishing a foundation for future studies of the circuit mechanisms underlying these diverse cerebellar functions.

## ACKNOWLEDGMENTS

This work was supported by R01 MH128744 to D.F.

